# Minimal-invasive 3D laser printing of microimplants *in organismo*

**DOI:** 10.1101/2024.01.23.576808

**Authors:** Cassian Afting, Philipp Mainik, Clara Vazquez-Martel, Tobias Abele, Verena Kaul, Kerstin Göpfrich, Steffen Lemke, Eva Blasco, Joachim Wittbrodt

**Affiliations:** Centre for Organismal Studies Heidelberg (COS), Heidelberg University, 69120 Heidelberg, Germany; Heidelberg International Biosciences Graduate School HBIGS, Heidelberg, Germany; HeiKa Graduate School on “Functional Materials”, Heidelberg, Germany; Institute for Molecular Systems Engineering and Advanced Materials (IMSEAM), Heidelberg University, 69120 Heidelberg, Germany; Organic Chemistry Institute (OCI), Heidelberg University, 69120 Heidelberg, Germany; Zentrum für Molekulare Biologie der Universität Heidelberg (ZMBH), Heidelberg University, 69120 Heidelberg, Germany; Max Planck Institute for Medical Research, 69120 Heidelberg, Germany; Institute of Biology, University of Hohenheim, 70599 Stuttgart, Germany

**Keywords:** additive manufacturing, 3D laser printing, two-photon lithography, microimplants, bioengineering, *Oryzias latipes*, *Drosophila melanogaster*

## Abstract

Multi-photon 3D laser printing has gathered much attention in recent years as a means of manufacturing biocompatible scaffolds that can modify and guide cellular behavior *in vitro*. However, *in vivo* tissue engineering efforts have been limited so far to the implantation of beforehand 3D printed biocompatible scaffolds and *in vivo* bioprinting of tissue constructs from bioinks containing cells, biomolecules, and printable hydrogel formulations. Thus, a comprehensive 3D laser printing platform for *in vivo* and *in situ* manufacturing of microimplants raised from synthetic polymer-based inks is currently missing.

Here we present a platform for minimal-invasive manufacturing of microimplants directly in the organism by one-photon photopolymerization and multi-photon 3D laser printing. Employing a commercially available elastomeric ink giving rise to biocompatible synthetic polymer-based microimplants, we demonstrate first applicational examples of biological responses to *in situ* printed microimplants in the teleost fish *Oryzias latipes* and in embryos of the fruit fly *Drosophila melanogaster*. This provides a framework for future studies addressing the suitability of inks for *in vivo* 3D manufacturing. Our platform bears great potential for the direct engineering of the intricate microarchitectures in a variety of tissues in model organisms and beyond.

## Introduction

Multi-photon 3D laser printing has been gaining attention in recent years for offering 3D fabrication of objects on the micro- and nanoscale^1, 2, 3, 4^. This technology has evolved originally from applications in mainly technical fields such as optics and photonics, but has also found its way into the life sciences thanks to the development of biocompatible materials. Today, this method is established in the life sciences for the precise fabrication of biocompatible scaffolds with subcellular resolution and applied in single cell, organoid, and cultured tissue research^4, 5, 6, 7, 8, 9, 10, 11, 12, 13, 14, 15, 16, 17^. The biocompatible scaffolds can be prepared either by physically encapsulating cells in a photocurable hydrogel or by printing and successive development, *i.e.*, removing excessive ink prior to cell seeding.

Application of printed 3D biocompatible scaffolds as implants possess great potential in experimental clinical and biological fundamental research^4, 18, 19, 20^. For example, cell-free and biocompatible structures have been printed and surgically transplanted into the subretinal space of pig eyes^12^. Histological and immunohistochemical analysis of these eyes then suggested an infiltration of the transplanted scaffold by retinal pigmented epithelium and photoreceptors within 30 days. However, major limitations of such surgical micro-implantations remain: they are technically challenging, the procedure is highly invasive, and damages to the surrounding tissues are almost always inevitable.

A well-explored alternative for targeting *in vivo* tissue engineering is the injection of cell-free or cell-laden hydrogels or ink-based composites for direct polymerization within tissues. Although a variety of physical and chemical *in vivo* crosslinking strategies have been explored for injected liquids^21, 22, 23, 24, 25^, these approaches are so far lacking control over the shape of the construct in 3D.

To address this issue, light-based 3D printing methods have very recently been explored for spatially controlled photopolymerization. For example, Chen *et al.* have reported noninvasive light-based *in vivo* 3D bioprinting of a gelatin-based ink with digitally guided near-infrared photopolymerization via micromirrors^26^. Furthermore, *in vivo* multi-photon 3D laser printing was recently presented by Urciuolo *et al.* for the fabrication of cell-laden and microscaled hydrogel structures^8^. Therein, the authors injected a cell-laden photosensitive gelatin-based prepolymer in skin, brain, or muscle tissue of anesthetized alive mice and used a multi-photon confocal microscope for printing. Similar to other bioprinting approaches, these efforts were aiming at *in vivo* manufacturing of tissue constructs, replicating the biochemical and mechanical properties of their surrounding tissues.

In a second work, *in vivo* 3D laser printing was performed outside of tissues, yet within living organisms. Conductive polypyrrole-containing microstructures were printed in the lumen of the *Caenorhabditis elegans* gut after feeding the worm with a hydrogel ink composition^5^. So far, these reports illustrated the potential of hydrogel-based ink systems in developed organisms.

Here, we present a platform for *in vivo, in organismo* manufacturing of microimplants by one-photon photocrosslinking and multi-photon 3D laser printing with potential in shaping tissue morphogenesis and eliciting biological response. For this, we employ a commercially available elastomeric ink, which allows us to manufacture biocompatible, synthetic polymer-based *micro*structures of low immunogenicity in alive developing organisms. With our approach, we have developed an *in vivo* tissue engineering toolbox for studying the effects of minimal-invasively delivered, 3D structured *micro*implants in (developing) organisms.

## Results

### Biocompatible one-photon photopolymerization in living, early organisms

The first step towards *in vivo* manufacturing of microimplants was to identify a suitable biocompatible material for *in vivo* one-photon photopolymerization. Towards this aim, we addressed photocrosslinkable elastomeric materials for *in vivo* biocompatibility of ink and photopolymerized material, *in vivo* ink stability and transparency. Not too low viscosity of the ink was a prerequisite for efficient microinjection, and eventually the microinjected material needed to be printable under *in vivo* conditions. A systematic analysis of inks indicated that the commercially available photocurable ink IP-PDMS (Nanoscribe GmbH, Germany) represented a very suitable material for this purpose. The ink is optically transparent with a refractive index of 1.45 (589 nm, 20°C) and upon photopolymerization gives rise to an elastomeric biocompatible material with a Young’s modulus of 15 MPa and polydimethylsiloxane-like properties^27^.

As *in vivo* model organisms, we selected the teleost fish *Oryzias latipes* (hereafter: medaka) and the fruit fly *Drosophila melanogaster* (hereafter: *Drosophila*). Their short generation time, their long history of genetic studies, their transparent eggs and the ease with which early developmental processes can be observed and experimentally addressed in these organisms place them both among the most important vertebrate and invertebrate models, respectively. Since their development occurs outside the mother and at sub-centimeter scales, all tissues are easily accessible via light irradiation during early development, which makes them uniquely suited to investigate the effects of tightly controlled *in vivo* polymerized materials on a wide range of different living tissues, organs and whole organisms.

To prepare these organisms for microinjection of ink and subsequent one-photon photopolymerization and multi-photon 3D laser printing, fertilized eggs were collected and then left to develop until they reached the required age. Fertilized *Drosophila* eggs were dechorionated immediately after deposition, mounted on a glass slide and covered with hydrocarbon oil. Fertilized medaka eggs were incubated overnight until embryonic stage 19-21^28^, the egg shell was then removed (dechorionation) and the embryos were mounted in custom made agarose molds. Following the microinjection of the ink, either into the yolk of developing *Drosophila* embryos or the developing optic vesicle of medaka embryos, embryos were exposed globally to a short 1 min UV light pulse to photocrosslink and thus solidify the microinjected ink in an one-photon photopolymerization process using the external light source of a conventional fluorescence microscope. In case of 3D laser printing, embryos were mounted for insertion into a Professional Photonic GT2 (Nanoscribe GmbH) and thus made available for 3D printing of microstructures. Finally, embryos were re-incubated for subsequent microscopic analysis (Fig. 1, Fig. 2a, c).

**Fig. 1:**
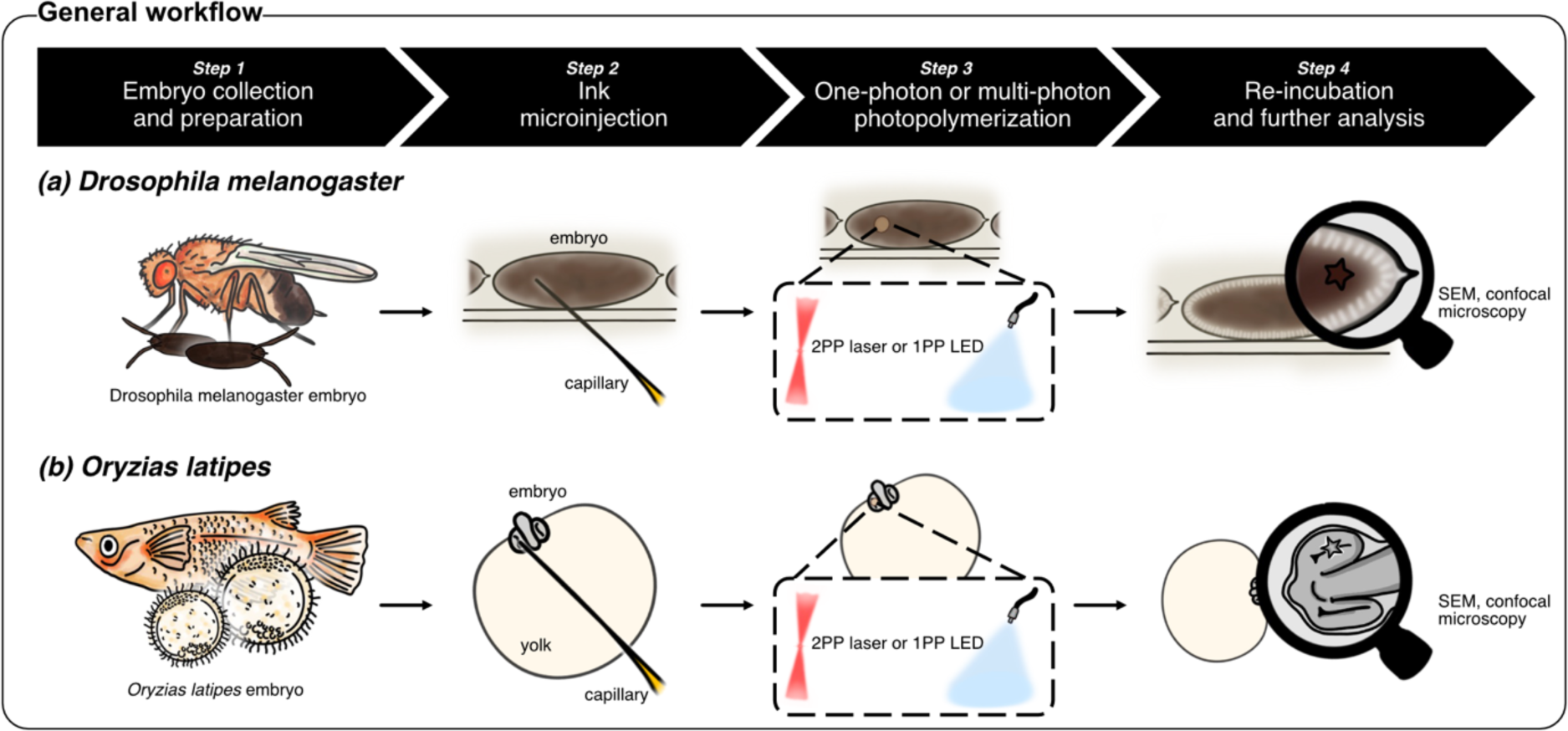
Biocompatible one-photon photopolymerization and multi-photon 3D laser printing workflow in living, early organisms. Schematic illustration of the one-photon photopolymerization (1PP) and multi-photon 3D laser printing (2PP) workflow in early *Drosophila melanogaster* and *Oryzias latipes* embryos.

**Fig. 2:**
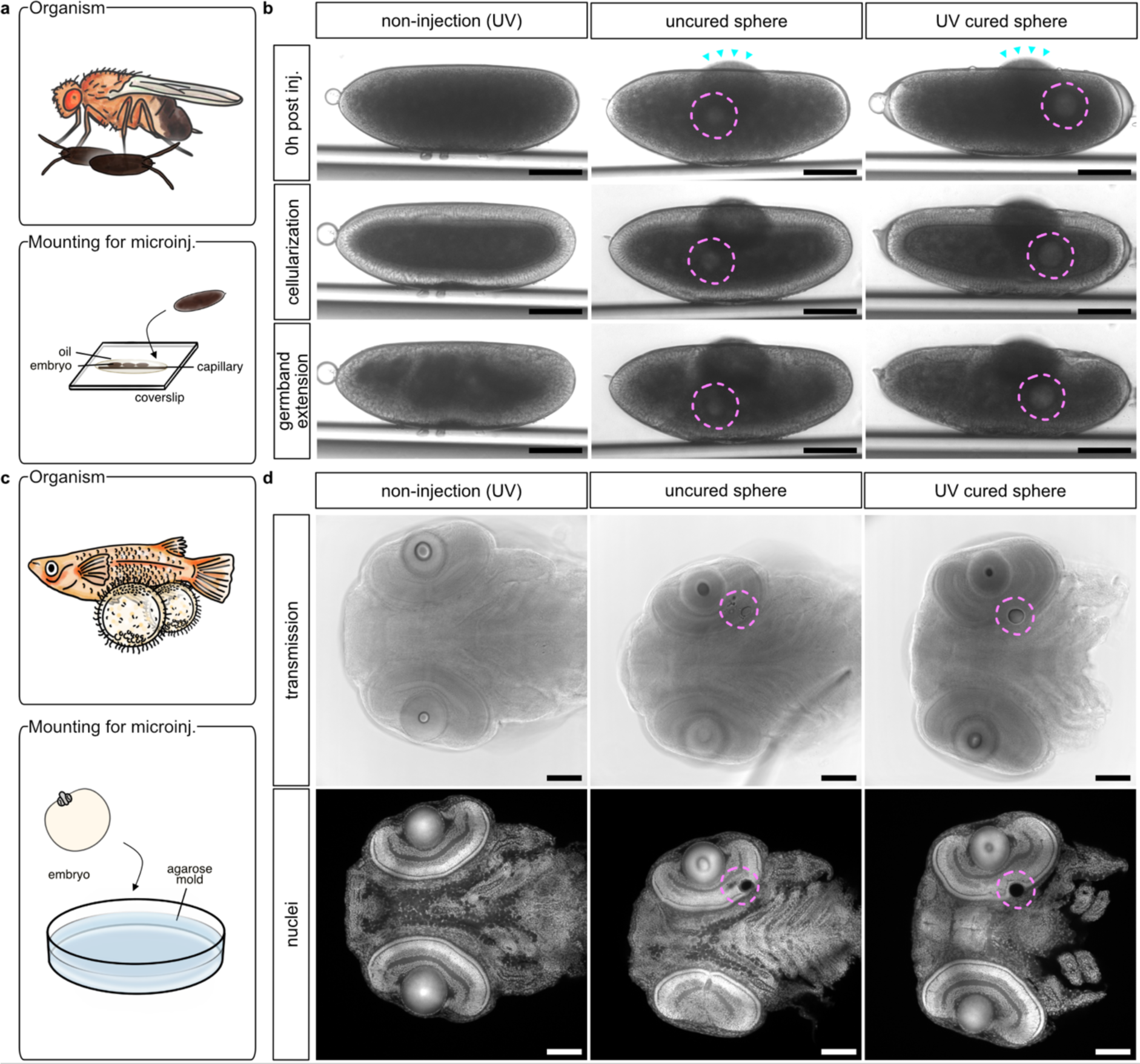
*In vivo* one-photon photopolymerization of microinjected ink does not adversely impact embryonic development. **a** Schematic illustration of the mounting method used to microinject ink in *Drosophila melanogaster* embryos. **b** Representative timelapse transmission microscopy of early *Drosophila melanogaster* embryos shortly after IP-PDMS microinjection (top), and during cellularization stage (middle) at about 2.5 hours post microinjection (hpi), and at about 3 hpi during germband extension stage (bottom). Microinjected IP-PDMS deposits were either left uncured (uncured sphere) or cured by UV light exposure (UV cured sphere). Non-injected controls (non-injection (UV)) were subjected to the same UV light regime. Because IP-PDMS is highly viscous, microinjection needles with a tip diameter of approx. 5-10 µm were used. During microinjection, slight leakages of yolk could be observed (indicated by cyan arrowheads), which however sealed off quickly and did not affect subsequent embryonic development. **c** Schematic illustration of the mounting method used to microinject ink in *Oryzias latipes* embryos. **d** Representative confocal microscopy of chemically fixed and whole-mount nuclear stained (Nuclei; DAPI) medaka embryo heads at 6 days post fertilization (dpf) after microinjection of IP-PDMS into their optic vesicles at 1 dpf. Dashed lines (magenta) indicate positions of microinjected deposits. Scale bars: 100 µm.

After microinjection into either the semi-aqueous extracellular environment of the yolk of early *Drosophila* embryos (1 hour post fertilization; 1 hpf) or the aqueous extracellular environment of the developing retina of medaka embryos (1 day post fertilization; 1 dpf), the IP-PDMS droplets adopted a stable and spherical shape. Even after microinjection into embryos, IP-PDMS photopolymerized efficiently (Fig. S1 showing UV cured spheres explanted from *Drosophila* embryos). Importantly, there was no indication of acute toxicity of IP-PDMS on the embryos as apparent by their unaffected development, the absence of immediate deformation, shrinkage or dissolution of ink surrounding tissue regardless of the microinjected IP-PDMS deposits being UV cured or left uncured.

For *Drosophila* embryos, timelapse transmission microscopy revealed that the embryo development progressed normally through all developmental stages followed in this study (cellularization and gastrulation) both post-microinjection and -photopolymerization. To control for UV toxicity during the polymerization process, non-injected controls were exposed to the same UV light dose and likewise showed normal gross development (Fig. 2b). For medaka embryos, normal development was confirmed by confocal microscopy of whole-mount nuclei stained heads at 6 dpf (5 days post-microinjection) with and without its UV curing, respectively. Notably, uncured IP-PDMS droplets consistently reduced in volume compared to UV cured spheres, suggesting a slow *in vivo* resorption of IP-PDMS ink over time (Fig. 2d).

### *In vivo* one-photon photopolymerized IP-PDMS permits targeted re-shaping of tissue morphogenesis

Following the establishment of microinjection and imaging conditions, we next analyzed the displacement of uncured and UV cured IP-PDMS spheres in *Drosophila* embryos during the process of gastrulation. Specifically, we asked whether overall gastrulation dynamics within the confined egg space affected uncured and UV cured spheres differently. For this, we microinjected IP-PDMS at the anterior and posterior poles of the embryos and tracked its total movement starting with the end of cellularization in 10 min increments for 2h in timelapse recordings acquired by transmission microscopy. We focused our analysis on embryos that had IP-PDMS spheres either in the anterior or in the posterior third of the embryo at the starting point of analysis (Fig. 3a). The total distance traveled of uncured and UV cured spheres was significantly increased if microinjected at the posterior pole compared to the anterior pole, but did not differ depending on their curation status (Fig. 3b). By contrast, if spheres were tracked only until the onset of gastrulation dynamics (i.e., from the last mitotic wave until the end of cellularization), no difference in total distance traveled between posteriorly and anteriorly microinjected deposits could be detected (Fig. S2). Taken together, these results suggest that increased displacement of posterior spheres during *Drosophila* gastrulation dynamics was driven mainly by the process of germ band extension, which pushes the posterior end of the embryo towards the anterior^29^.

**Fig. 3:**
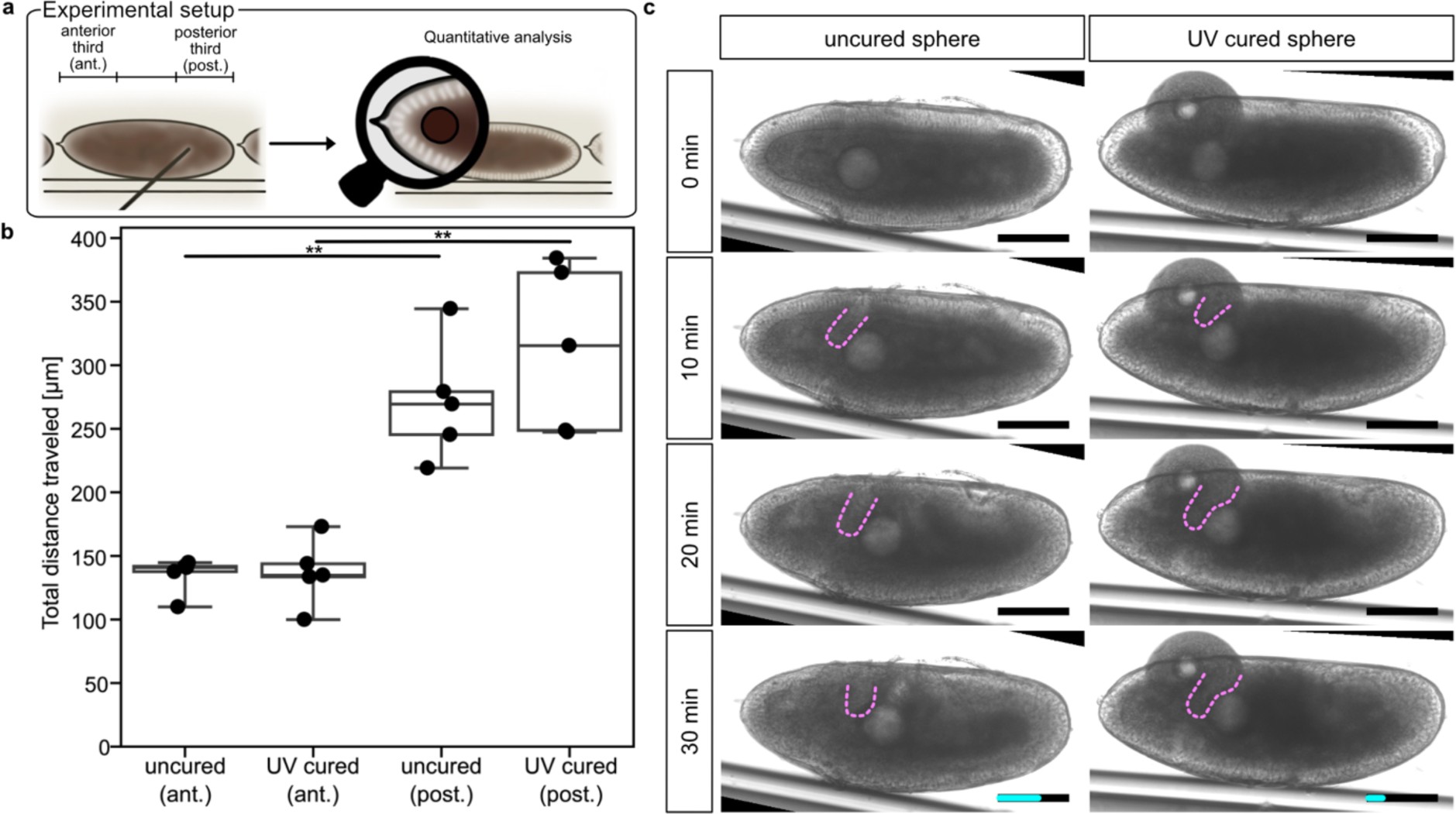
*In vivo* one-photon photopolymerized IP-PDMS spheres allow local re-shaping of tissue morphogenesis. **a** Schematic illustration of microinjection scheme for quantitative analysis of uncured and UV cured IP-PDMS deposit dynamics in *Drosophila melanogaster (Drosophila)* embryos. **b** Quantification of total distance traveled of the uncured and UV cured, microinjected IP-PDMS deposits at the anterior (ant.) and posterior (post.) poles of the *Drosophila* embryos. Spheres were tracked in 10 min increments in timelapse transmission images from the end of cellularization 2h onwards. Boxplots include data from 5 embryos each with boxes indicating 25-75% percentiles and whiskers 10-90% percentiles. Individual data points shown as dots. Student’s *t*-test with unequal variance; **p-value: 0.002. **c** Representative timelapse transmission microscopy of early *Drosophila* embryos microinjected with IP-PDMS during cephalic furrow formation. Microinjected IP-PDMS deposits were either left uncured (left panels) or cured by UV light exposure (right panels). Magenta dashed lines indicate the gross morphology of the cephalic furrow, cyan scale bar insets indicate the total distance traveled of the spheres within 15 min after the first contact with the emerging cephalic furrow. Scale bars: 100 µm.

While the overall *in vivo* movement dynamics of uncured and UV cured IP-PDMS spheres in *Drosophila* embryos did not differ, we could observe that UV cured spheres had a striking capacity to alter local tissue dynamics, which was not seen with uncured spheres. This could be demonstrated exemplary for a transient epithelial fold at the head/trunk interface, the so-called cephalic furrow (CF). While the dynamics CF formation remained unaffected in the presence of uncured spheres, a UV cured sphere has the capacity to alter the shape of the cephalic furrow upon contact. Consistent with the notion that such capacity to change the shape of an epithelial fold was associated with direct sphere contact, we found that the UV cured sphere traveled less than half the distance compared to the uncured droplet within 15 min upon contact with the CF (Fig. 3c).

### Biocompatible multi-photon 3D laser printing in living, early organisms

In addition to the described one-photon photopolymerization process, IP-PDMS can be shaped with higher precision and to complex geometries using multi-photon 3D laser printing. Therefore, we adapted the workflow to 3D laser print inside of the uncured IP-PDMS droplet in living, early organisms using a Professional Photonic GT2 (Nanoscribe GmbH). For *Drosophila* embryos, we did not need to modify the one-photon polymerization pipeline to enable insertion into the substrate holder of the 3D laser printer and subsequent 3D printing (Fig. 4a), while for medaka embryos the mounting method had to be adapted. Here, insertion into the substrate holder and subsequent 3D printing was possible after immobilizing the embryo on the insertable cover glass in a low-melting point agarose dome following its microinjection with IP-PDMS. Orientation with the embryos’ heads down towards the cover glass proved pivotal to not exceed the working distance of the 25x objective (Fig.4c; see Methods for details).

**Fig. 4:**
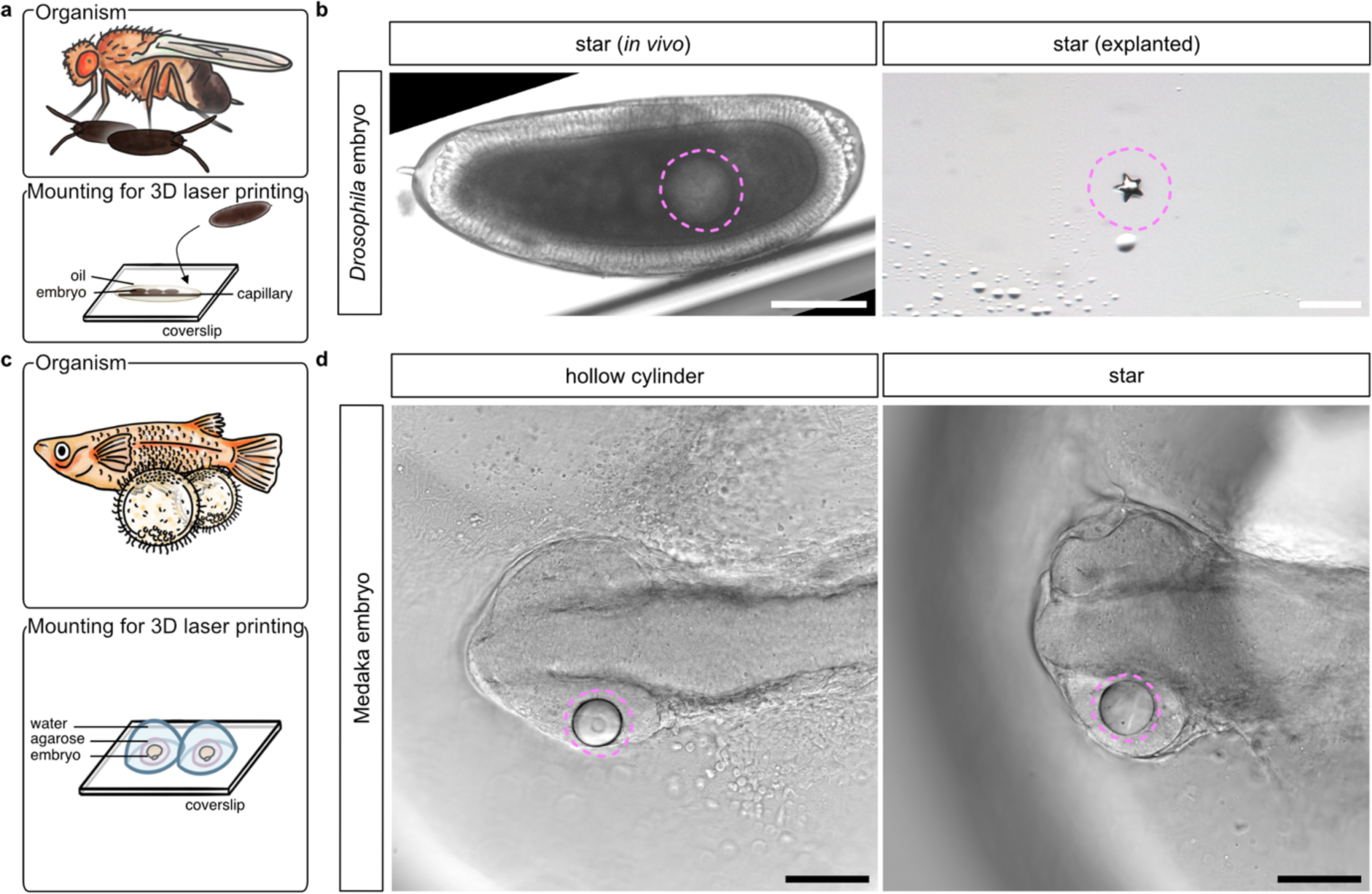
Microinjected ink deposits can be used for *in vivo* multi-photon 3D laser printing of microstructures. **a** Schematic illustration of the mounting method used to enable 3D laser printing in *Drosophila melanogaster (Drosophila)* embryos. **b** Representative transmission microscopy of early *Drosophila* embryos microinjected with IP-PDMS and subsequently subjected to 3D laser printing of a 40 µm x 40 µm x 13 µm star. The printed structure within the yolk of developing *Drosophila* embryos were then manually extracted by dissection and stereomicroscopically imaged. **c** Schematic illustration of the mounting method used to enable 3D laser printing in *Oryzias latipes* (medaka) embryos. **d** Representative transmission microscopy of stage 19-21 medaka embryos microinjected with IP-PDMS and subsequently subjected to multi-photon 3D laser printing of a 50 µm x 50 µm x 20 µm hollow cylinder (20µm inner diameter) and a 60 µm x 60 µm x 20 µm star. Dashed lines (magenta) indicate positions of microinjected deposits. Scale bars: 100 µm.

In this way we were able to print structures *in vivo* with sizes down to the low microscale with even smaller feature sizes. Timelapse transmission microscopy of *Drosophila* embryos with 3D laser printed stars (40 µm x 40 µm x 13 µm) showed that gross embryonic development is not perturbed by multi-photon 3D laser printing. The printed structures were not visible at any time in transmission mode, presumably due to the embryo’s thickness and the yolk’s optical properties. However, dissection of the embryos confirmed successful microprinting of the structures (Fig. 4b). Fig. 4d shows transmission microscopy of exemplary medaka embryos of different stages with different 3D printed microstructures in their developing retinas (from left to right: hollow cylinder 50 µm x 50 µm x 20 µm (20 µm inner diameter), star 60 µm x 60 µm x 20 µm).

### *In vivo* multi-photon 3D laser printed microstructures spontaneously develop and fully integrate into surrounding tissues without significant immunogenicity

After 3D laser printing, printed structures float in the unpolymerized ink within the embryos and are thus initially not directly contacting the cells of the adjacent tissue. In medaka embryos, structures remained floating in the ink residue for days as embryonic development continued. Nevertheless, raising medaka embryos after multi-photon 3D laser printing to the end of embryonic development at stage 41^28^ (18 days post microinjection, Fig. 5a) resulted in spontaneous and mostly complete removal of the unpolymerized ink in a strain-specific manner, presumably by slow resorption of the non-toxic IP-PDMS over time.

**Fig. 5:**
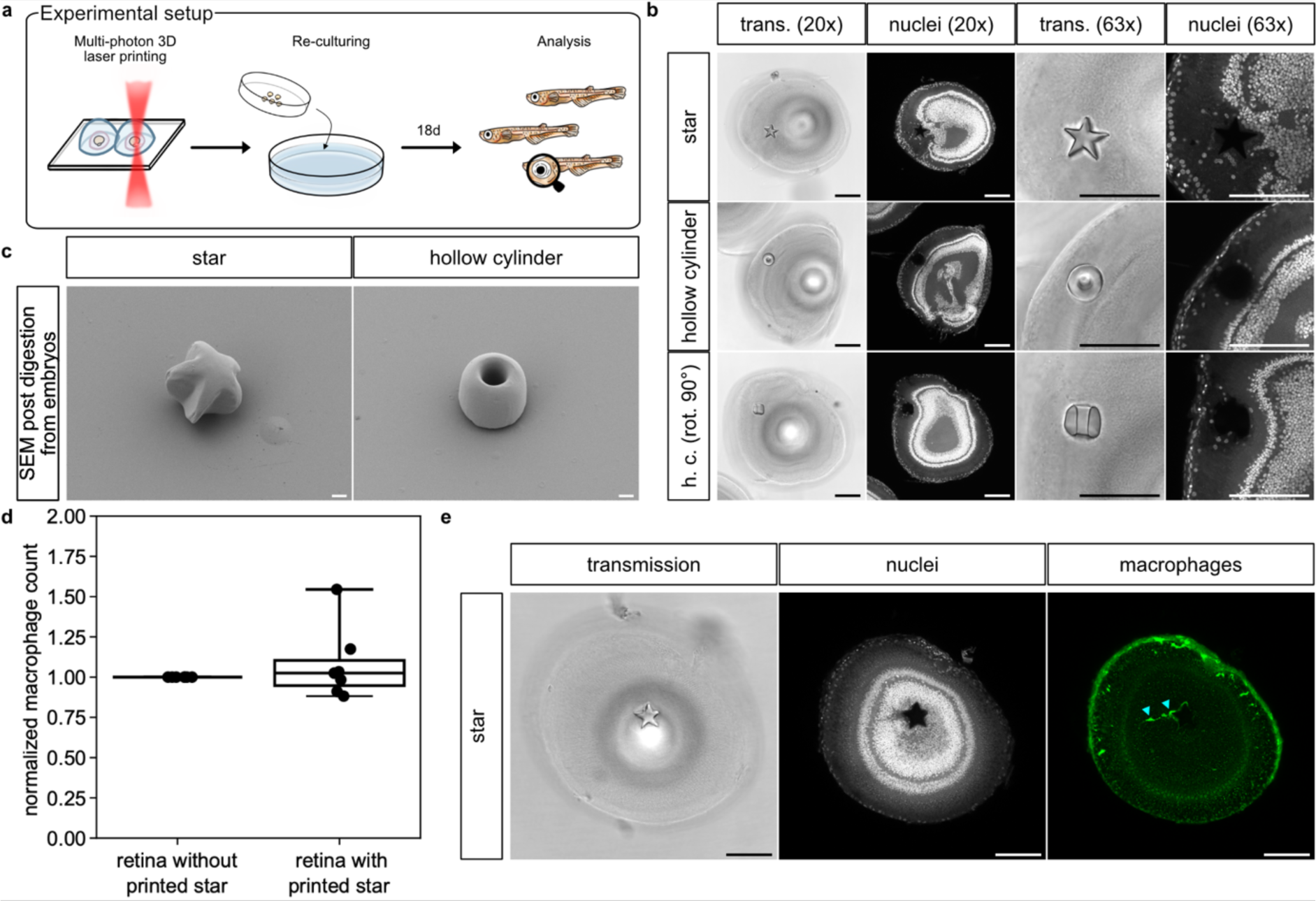
*In vivo* multi-photon 3D laser printed microstructures develop spontaneously in the developing medaka retina, fully integrate into the tissue and show no significant immunogenicity. **a** Schematic illustration of the experimental setup that led to the images shown in b. **b** Representative confocal microscopy of chemically fixed and whole-mount nuclear stained (Nuclei; DAPI) medaka embryo eyes (stage 41) after microinjection of IP-PDMS into their optic vesicles and subsequent 3D laser printing at 1 dpf. Printed structures measure 60 µm x 60 µm x 20 µm for the star and 50 µm x 50 µm x 20 µm for the hollow cylinder (20 µm inner diameter; shown in two orientations). Scale bars: 100 µm. **c** Scanning electron microscopy of microprinted structures after enzymatic extraction from medaka embryo eyes obtained like shown in a. Scale bars: 10 µm. **d** Quantification of total macrophage count in the whole retinas of n = 7 embryos with printed stars (60 µm x 60 µm x 20 µm). Cell counts were normalized to the cell counts of the contralateral control retinas of the same embryo without printed stars to account for differential cxcr3a::GFP expression levels between individual embryos. Boxplot boxes indicating 25-75% percentiles with whiskers indicating 10-90% percentiles. Individual data points shown as dots. **e** Representative confocal microscopy of chemically fixed and whole-mount nuclear and anti-GFP stained (Nuclei: DAPI; Macrophages: Cxcr3a::GFP) medaka embryo retina (stage 41) after microinjection of IP-PDMS into its optic vesicle and subsequent 3D laser printing at 1 dpf. Printed star measures 60 µm x 60 µm x 20 µm. Cyan arrowheads indicate two individual macrophages in close proximity to the star. Scale bars: 100 µm.

Whole-mount nuclear stain and imaging by confocal microscopy revealed an integration of the structures into the surrounding tissue, most frequently into the retinal outer segments layer (OSL), seamlessly without any apparent signs of immunogenic reactions, scarring, or developmental defects. Moreover, multiple nuclei consistently assembled around the structures, locally breaking the otherwise cell soma free organization of the OSL. This was observed independently of the shape of the printed structures and their position within the retinal layering since nuclei with morphologies different from surrounding cells were observed to neighbor printed structures intermittently even in other retinal layers like the retinal inner nuclear layer (INL; Fig. S3a). Cells did not actively invade the hollow cylinders with openings 20 µm in diameter and rather stayed in closer connection to cells of the surrounding tissue (Fig. 5b; contralateral control eyes in Fig. S3b). Scanning electron microscopy of printed structures enzymatically extracted from stage 41 embryonic medaka retinas revealed a remarkable printing precision given the particular printing environment, albeit showing a tendency towards round distortions if printed close to the edge of the ink bubble likely due to optical diffraction (Fig. 5c, 3D rendered models of microstructures in Fig. S4).

To address the apparent absence of a significant immunogenic reaction to the microstructures, we employed a transgenic reporter line labeling macrophages (cxcr3a::GFP), the known key players in the immediate to long-term immune response to implanted biomaterials^30, 31, 32^ and compared the total number of macrophages in the experimental and control retina. At 18 days post ink microinjection and 3D laser printing we consistently found comparable macrophage numbers in both retinas, with no significant difference when comparing retinas with microstructures to contralateral control retinas without (Fig. 5d). This indicates that there was no immunogenic response triggering the infiltration of additional macrophages in the microinjected/printed retinas as a chronic inflammatory response. Moreover, we consistently observed a macrophage population of a few cells at the sites of the microstructures inspecting the 3D printed object during immunosurveillance without triggering an adverse immunogenic reaction (Fig. 5e). The cytosolic fluorescent signal of labeled macrophages was found to partially overlap with the above-mentioned nuclei.

## Discussion

In this work, we outline a platform for one-photon photopolymerization and also adapted it to enable multi-photon 3D laser printing in early alive *Drosophila* and medaka embryos. We show how one- and multi-photon polymerizable polydimethylsiloxane can be microinjected, and UV cured as well as 3D printed in their extracellular tissue environments. Regarding requirements for *in vivo* printable synthetic polymers, we found it to be crucial for the biocompatible ink to be hydrophobic to form stable droplets with a low solubility of the different ink components in aqueous environments. Furthermore, the ink droplets need to keep their integrity and printability between microinjection and UV curing or 3D laser printing.

In case of *in vivo* one-photon photopolymerization, we observed that only after UV induced photocrosslinking, the PDMS-like material could locally re-shape tissue morphogenesis in *Drosophila* embryos in ways that have not been possible by established methodology. This might be the consequence of an interaction of the basal cell side with the UV cured IP-PDMS sphere, which the fluid surface interface of uncured IP-PDMS might not permit. A non-natural point of adhesive interaction of the yolk-facing cells in the emerging cephalic furrow might then be an anchor point for the tissue to re-shape around. PDMS is generally known to be biocompatible but unfavorable for cell adhesion. Coating with extracellular matrix (ECM) proteins via physisorption is regularly used to increase cell attachment on PDMS surfaces^33,34^. Specifically for multi-photon printed IP-PDMS, it has recently been shown to be able to support cell attachment of human mesenchymal stem cells after fibronectin functionalization^9^. Given that our PDMS-like material microstructures reside outside bona fide epithelia of the *Drosophila* and medaka embryos, a natural coating with ECM or yolk resident proteins via physisorption after UV curation or 3D printing seems likely. On top of that, the PDMS-like material structures are being exposed to the composition and spatial distribution of ECM and yolk resident proteins that are specific to the embryo, its developmental stage and the tissue in question and therefore likely suitable for supporting attachment of cells of that particular tissue.

In late medaka embryos we found the 3D microstructures to develop and subsequently integrate into the surrounding tissue in less than 3 weeks after ink microinjection and 3D laser printing. Depending on the specific position of the 3D microstructures within the retinal layers, they were found juxtaposed to the outer segments of the photoreceptors in the OSL or the cell somata of the cells in the INL, permitting direct interaction with the cells of the tissue. This opens up the tissue microarchitecture to precise *in vivo* tissue engineering efforts utilizing the physicochemical features of a given printable biomaterial. Here, among others, properties like stiffness could be exploited as stiffness has been widely characterized to be a critical physical stem cell niche parameter and key determinant of differentiation^35, 36^, proliferation^37^ and motility^38^.

Moreover, we found IP-PDMS microstructures to have a remarkably low immunogenicity. At 18 days post ink microinjection and 3D laser printing, no macrophage infiltration of the retina was observed, showing that the *in situ* manufactured microstructures did not trigger an immunogenic response. Few resident macrophages were found to partially surround the printed microstructures with their cell bodies and pseudopodia, reminiscent of a recognition of the structure during immunosurveillance without them triggering an adverse immunogenic reaction to the foreign body. Inflammatory responses to immunogenic events, like an injury, generally are known to trigger massive infiltration of macrophages into the organ in question and to the site of injury even in early and late fish embryos^39, 40^. The lack of an inflammatory response reflects the high degree of biocompatibility of the material as well as the minimal-invasive nature of the presented platform, since the tissue damage during surgical implantation releases damage-associated molecular patterns (DAMPs) and thus always results in an inflammatory response^31, 32^. *In vivo* biocompatibility studies following mainly subcutaneous, intramuscular and intraperitoneal implantation protocols for PDMS-based prosthetic macroscopic implants showed a high degree of biocompatibility with only mild acute and chronic inflammatory responses in the past^41^.

In conclusion, this study presents the first account of *in vivo* fabrication of 3D artificial microimplants inside the organ of a developing organism, using both one-photon and multi-photon polymerization. Importantly, this approach offers a framework for the minimal-invasive installation of biocompatible and non-degradable microimplants *in vivo* and thus provides opportunity to study and apply the *in vivo* bioengineering potential of materials and their shapes in 3D within tissue microarchitectures. Our pipeline allows for studying long-term interactions of complex *in vivo* tissue environments on micromaterials and their 3D integrity and *vice versa*. We identified an easily accessible, biocompatible, *in vivo* printable material of particularly low immunogenicity that is capable of mediating biological responses in extracellular *in vivo* environments. It is enticing to speculate how the presented platform is a first step towards *in vivo* 3D manufacturing of microimplants to treat human diseases affecting tissue or organ microarchitectures.

## Methods

### Materials

IP-PDMS (IP-polydimethylsiloxane; Nanoscribe GmbH, Germany) was used as the microinjectable, biocompatible ink in this study. The ink was protected from UV light prior to and after microinjection and mostly handled under yellow light conditions.

### Fish husbandry and maintenance

Medaka fish (*Oryzias latipes*) stocks were maintained according to the local animal welfare standards (Tierschutzgesetz §11, Abs. 1, Nr. 1, husbandry permit AZ35-9185.64/BH, line generation permit number 35–9185.81/G-145/15 Wittbrodt). The fish are being kept as closed stocks in constantly recirculating systems at 28°C with a 14h light/10h dark cycle. The following medaka lines were used in this study: Heino strain as an albinism exhibiting mutant^42^ and cxcr3a::GFP^43^.

### Fly line and embryo handling

*OregonR* line was used as wildtype *Drosophila melanogaster*. Embryos were collected on apple juice containing agar plates (1:4 mix) with yeast paste at room temperature and subsequently prepared for microinjection. Embryos were dechorionated using bleach for 45s and afterwards washed thoroughly with tap water. Dechorionated embryos were transferred on a 22 x 22 mm cover glass, aligned against a glass capillary and dried at room temperature for 7.5 min for slight volume reduction. Embryos were finally covered with a mixture of hydrocarbon oil (700 and 27 at a 4:1 ratio; Sigma Aldrich) and subjected to microinjection in less than 1h post fertilization. One-photon photopolymerization, multi-photon 3D laser printing and further microscopic analysis were performed typically within an hour of microinjection with embryos left mounted onto the cover glass.

### Fish embryo handling

Medaka embryos were collected at day 0 shortly after fertilization and incubated in embryo rearing medium (ERM; 17 mM NaCl, 40 mM KCl, 0.27 mM CaCl_2_, 0.66 mM MgSO_4_, 17 mM HEPES) at temperatures between 18°C and 32°C depending on the desired timing of the target stage. Stage 19-21^28^ (1 day post fertilization; dpf) embryos were subjected to dechorionation using hatching enzyme, washed and kept in 100 U/ml penicillin-streptomycin (P/S) containing ERM. Embryos were transferred into 1% agarose molds commonly used for transplantations^44^, oriented heads down for microinjection and punctured at the vegetal pole. Microinjected embryos to be subjected to UV curation of the microinjected ink were transferred into 100µl of P/S containing ERM. For multi-photon 3D laser printing, embryos were imbedded in 1% low-melting point agarose (Carl Roth, Cat#: 6351.5) solved in ERM onto a 22 x 22 mm cover glass and oriented with the heads towards the glass, minimizing the distance to the printer’s objective. UV curation and 3D laser printing were conducted within an hour of microinjection. Low-melting point agarose domes were moisturized with ERM and inserted into the Photonic Professional GT2 sample holder. After printing, embryos were manually extracted from the agarose using tweezers. Finally, embryos were re-incubated on glass ware in 100 U/ml P/S containing ERM until either 6 dpf or hatchling stage (s41^28^, 19 dpf) with daily assessment of their gross morphology by stereomicroscopy. In order to prevent unwanted UV curation of unprinted IP-PDMS within the embryos, embryos were raised in total darkness until stage s40^28^.

### Microinjection of embryos

For microinjection, borosilicate micropipettes (1 mm OD x 0.58 mm ID x 100 mm L; Warner Instruments, Cat#: 30-0016) were pulled on a Flaming/Brown micropipette puller P-97 (Sutter instruments Co.) with the following settings: Heat 505, Pull 25, Velocity 250, Time 10, 1 cycle to create pulled pipettes with a slender taper. Pulled micropipettes were cut open manually to have a small, beveled opening of around 5 to 10 microns. Microinjections were performed with either a CellTram 4m oil microinjector (Eppendorf AG) or homemade oil microinjector and a standard manual micromanipulator under an epifluorescence stereomicroscope (Olympus MVX10; MV PLAPO 1x objective).

### One-photon photocrosslinking

For one-photon photocrosslinking of the ink (UV curing) medaka embryos kept in 100µl of P/S containing ERM or *Drosophila* embryos mounted on the cover glass were exposed for 60 s at a 1 cm distance to unfiltered Leica EL6000 light (100% intensity; Lamp: HXP-R120W/45C VIS, power input 120W, Osram Licht AG, Munich, Germany). Successful solidification was evaluated for every individual experiment by dissection of the embryos, removal of the UV cured sphere and manual crushing or drying of it.

### Multi-photon 3D laser printing

Multi-photon 3D laser printing was performed with the commercial 3D printing system Photonic Professional GT2 (Nanoscribe GmbH) with a laser wavelength of 780 nm and a 25x objective (NA=0.8). STL files of desired 3D structures were transformed into Nanoscribe GWL printing files using the Describe software. In this software, slicing and hatching were set to 300 nm for all geometries, as well as galvo and piezo modes were selected for xy and z movement of the laser, respectively.

Before printing in the microinjected droplet, precise laser positioning was required in *xyz* dimensions. For this purpose, a glass slide with pure IP-PDMS was first prepared for printing in oil immersion mode with Immersol 518 F from Carl Zeiss AG. Subsequently, simple crosshair structures were printed to align the *xy* position of the laser to an external crosshair in the camera. Afterwards, the glass slide was removed from the substrate holder and replaced by the glass slides with the embryos (medaka and *Drosophila*).

The first initial *xy* positioning was followed by preparing the slides for printing in oil immersion mode and inserting the equipped substrate holder in the printer. The ink droplet in the embryo was positioned in the center of the external camera crosshair for *xy* positioning. Now, moving of the stage (in *z* axis) was used to position the droplet in the focus plane. After final *xyz* positioning, 3D laser printing of desired 3D microstructures was performed with a scan speed of 20 mm/s and laser power of 30 mW. After printing, the oil on the bottom of the glass slides was removed with HPLC grade isopropanol.

### Fly embryo live imaging

Following either one-photon photocrosslinking or multi-photon 3D laser printing, cover glasses with *Drosophila* embryos mounted on were adhered onto standard glass microscopy slides. Live embryos were timelapse imaged at a framerate of 1 frame every 1 min over the course of 7 h, resulting in videos capturing their gross development after intervention. 633 nm laser powered transmission images were acquired on a Leica SP5 DMI6000CS inverted confocal microscope using the 10x objective with 2.5x digital zoom. Microinjected deposits were tracked using manual ROI specification in ImageJ^45^.

### Fluorescent labeling and fish embryo imaging

For whole-mount fluorescent staining, embryos were fixed overnight in 4% PFA at 4°C and washed with PTW (0.05% Tween20 solved in PBS). Either full heads, eyes or retinas were manually dissected from the embryos using tweezers. Dissected samples were bleached (either 0.3% (Heinostrain) or 3% (cxcr3a::GFP) H_2_O_2_, 0.5% KOH; dissolved in PTW) for 15-30 min at room temperature and washed 5 times with PTW. Sample permeabilization was performed with acetone for 15 min at −20°C. Samples were blocked with 4% sheep serum, 1% BSA and 1% DMSO in PTW for 1 h at room temperature. Nuclear stain was carried out overnight at 4°C with DAPI (10 µg/ml in DMSO, Carl Roth, Cat#: 6335.1) solved in PTW. Cxcr3a::GFP samples were incubated with primary antibodies against GFP for 48 h at 4°C (chicken anti-GFP (Thermo Fisher Scientific, Cat#: A10262; 1:300)). Samples were washed five times with PTW and subsequently incubated with the respective secondary antibodies (1:500) for 24 h at 4°C (donkey anti-chicken Alexa Fluor 488 (Jackson ImmunoResearch Europe Ltd., Cat#: 703-545-155)). In preparation for imaging, embryos were washed five times in PTW and finally transferred into optical clearing solution^46^ (20% (wt/vol) urea (Sigma Aldrich, Cat#: V900119), 30% (wt/vol) D-sorbitol (Sigma Aldrich, Cat#: V900390), 5% (wt/vol) glycerol (Sigma Aldrich, Cat#: V900122) dissolved in DMSO (Sigma Aldrich, Cat#: V900090)). Imaging was performed on a Leica TCS Sp8 inverted confocal microscope (20x and 63x oil immersion objective).

Macrophages were manually counted in whole retinas using the Cell Counter plugin in ImageJ^45^.

### Scanning electron microscopy (SEM)

3D laser printed structures were extracted from fixed medaka embryo samples via overnight incubation at 60°C in extraction buffer (100 nM Tris-HCl pH 8.5, 10 mM EDTA, 200 mM NaCl, 1% SDS, 1 mg/ml Proteinase K (Sigma Aldrich, Cat#: 3115852001)), then washed with isopropanol, and transferred onto glass coverslips. The glass coverslips containing the 3D structures were then sputter coated with a 12 nm layer of Pt/Pd (80:20) and imaged using a field-emission scanning electron microscope (Ultra 55, Carl Zeiss Microscopy) operated at a primary electron energy of 3 keV with the chamber SE2 detector and a tilting angle of 30°.

### Statistical analysis

Two-tailed Student’s *t*-test with unequal variance was used for the calculation of significant differences. Differences between two groups with p-values < 0.05 were considered statistically significant.

## Acknowledgements

The authors thank Dr. Girish Kale for help with *Drosophila melanogaster* timelapse imaging analysis and Maily Scorcelletti for help with setting up *Drosophila melanogaster* experiments. The authors thank Prof. Dr. Rasmus Schröder (Heidelberg University) for the access to the electron microscopy facilities. C.A. was partially funded by the Structured Doctoral programme (*Strukturiertes Doktorandenprogramm zum Erwerb des Dr. med. und Dr. rer. nat.*) of Heidelberg University. K.G., E.B. and J.W. acknowledge the funding from the Excellence Cluster “3D Matter Made to Order” (EXC-2082/1-390761711). E.B. further acknowledges funding from the Carl Zeiss Foundation through the “Carl-Zeiss-Foundation-Focus@HEiKA”.

C.V.M. acknowledges the Fonds der Chemischen Industrie for the support during her PhD studies through the Kekulé Fellowship.

## Author contributions

Conceptualization: C.A., P.M., E.B., J.W.

Investigation: C.A., P.M.,C.V.M.,T.A., V.K., K.G., S.L., E.B., J.W.

Visualization: C.A., P.M.,C.V.M.

Funding acquisition: S.L., K.G., E.B., J.W.

Project administration: E.B., J.W.

Supervision: S.L., K.G., E.B., J.W.

Writing - original draft: C.A., P.M., E.B., J.W.

