## Supplemental material for "Minimal-invasive 3D laser printing of microimplants *in organismo*"

**Contains Figure S1-S4**

**Fig. S1: UV cured spheres explanted from *Drosophila* embryos after timelapse microscopy.**

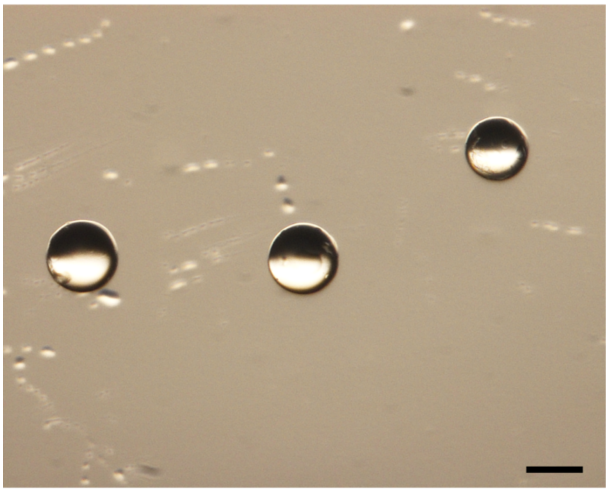

Scale bar: 100 µm.

**Fig. S2: Total distance traveled of uncured and UV cured spheres before cellularization and images without overlays shown in Figure 3.**

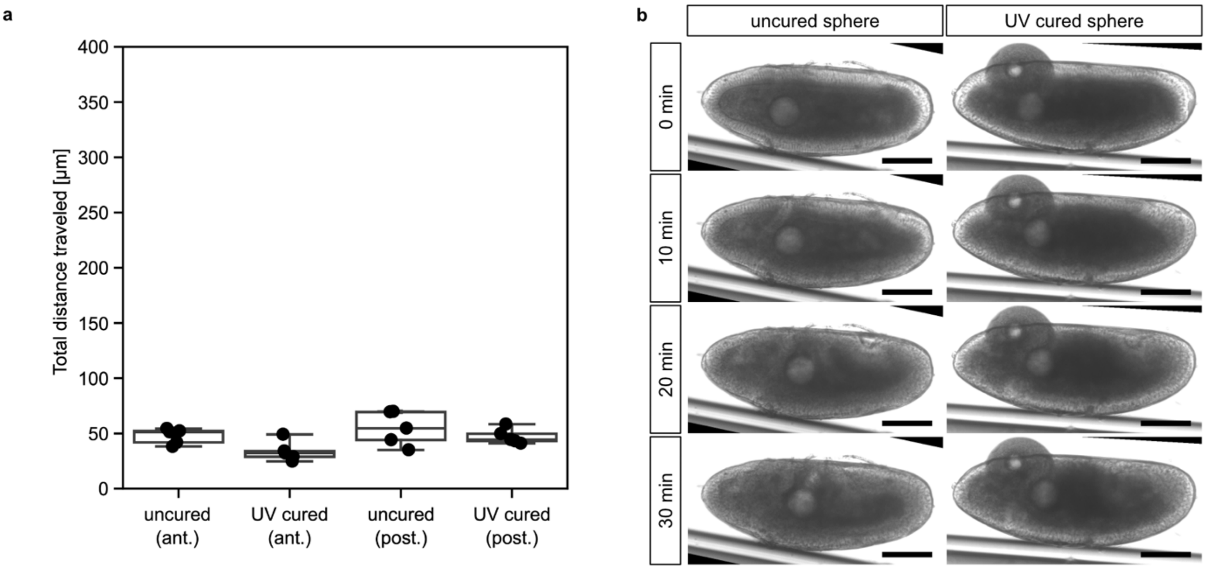


**a** Quantification of total distance traveled of the uncured and UV cured, microinjected IP-PDMS deposits at the anterior (ant.) and posterior (post.) poles of the *D. melanogaster* embryos. Spheres were tracked in 10 min increments in timelapse microscopy images from the last mitotic wave until end of cellularization. Boxplots include data from 5 embryos each with boxes indicating 25-75% percentiles and whiskers 10-90% percentiles. Individual data points shown as dots. **b** Corresponding images without overlays to images shown in Figure 3c. Scale bars: 100 µm.

**Fig. S3: Multi-photon 3D laser printed star in bipolar cell layer and contralateral control eyes of the stage 41 medaka embryos shown in Figure 5.**

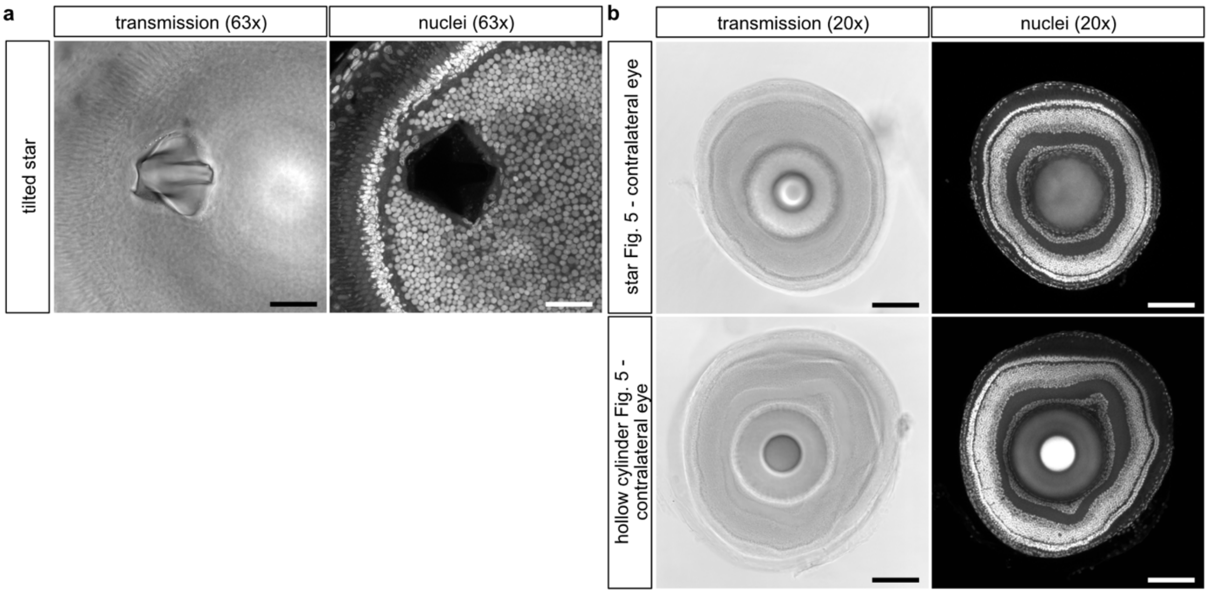


**a** Representative confocal microscopy of chemically fixed and whole-mount nuclear stained (Nuclei; DAPI) medaka embryo eyes at 19 days post fertilization (dpf) after microinjection of IP-PDMS into their optic vesicles and subsequent 3D laser printing at 1dpf. The slightly tilted star measures 60 µm x 60 µm x 20 µm and is integrated into the retinal inner nuclear cell layer (INL). **b** Contralateral, uninjected eyes of the medaka embryos shown in Figure 5b. Scale bars: 100 µm.

**Fig. S4: 3D rendered models of in vivo 3D printed microstructures shown in Fig. 4 and 5.**

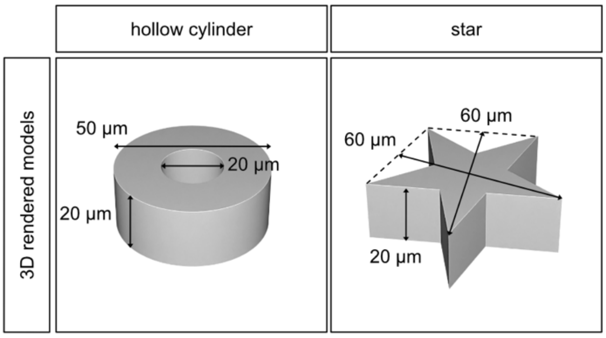
